# Neural synchrony predicts children’s learning of novel words

**DOI:** 10.1101/2020.07.28.216663

**Authors:** Elise A. Piazza, Ariella Cohen, Juliana Trach, Casey Lew-Williams

**Affiliations:** Department of Psychology, Princeton University; Princeton Neuroscience Institute, Princeton University; Department of Brain & Cognitive Sciences, University of Rochester; Icahn School of Medicine at Mount Sinai; Department of Psychology, Yale University

**Keywords:** word learning, book reading, neural synchrony, fNIRS

## Abstract

Social interactions have a well-studied influence on early development and language learning. Recent work has begun to investigate the neural mechanisms that underlie shared representations of input, documenting neural synchrony or coupling (measured using intersubject temporal correlations of neural activity) between individuals exposed to the same stimulus. Neural synchrony has been found to predict the quality of engagement with a stimulus and with communicative cues, but studies have yet to address how neural synchrony among children may relate to real-time learning. Using functional near-infrared spectroscopy (fNIRS), we recorded the neural activity of 45 children (3.5-4.5 years) during joint book reading with an adult experimenter. The custom children’s book contained four novel words and objects embedded in an unfolding story, as well as a range of narrative details about object functions and character roles. We observed synchronized neural activity between child participants during book reading and found a positive correlation between learning and intersubject neural synchronization in parietal cortex, an area implicated in narrative-level processing in adult research. Our findings suggest that children’s learning is facilitated by active neural engagement with the dynamics of natural social input.

## Introduction

Social interaction and communication are central in facilitating children’s early learning. Language learning, including the acquisition of lexical, syntactic, and pragmatic knowledge, is highly dependent on social engagement with adults and other children, which helps guide learners’ attention toward the most useful input (Tomasello, 1992). While recent work has begun to characterize the neural correlates of social communication in adults (Silbert et al., 2014; Stephens, Silbert, & Hasson, 2010), the neural mechanisms underlying learning and social engagement in young children are largely unknown. In this study, we recorded activity from children’s brains while they engaged in natural, joint book reading to investigate the neural mechanisms underlying learning during social communication with adults. We focused in particular on children’s learning of novel words embedded in a storybook.

If the primary function of language is to communicate ideas and intentions to others, it is therefore unsurprising that language learning relies on communicative contexts – an idea that has been investigated in a wealth of previous research. For example, in one study, infants learned sound categories better from in-person interactions than from audio or audiovisual recordings (Kuhl, Tsao, & Liu, 2003), and in another, infants only succeeded in learning a pattern embedded in a novel auditory signal if it was presented in a communicative context (Ferguson & Lew-Williams, 2016). In addition, contingent social feedback from caregivers, including smiling, touching, and vocalizing, has been shown to shape the acoustic features of infants’ babbling (Goldstein, King, & West, 2003; Goldstein & Schwade, 2008). The ability to engage in joint attention with adults emerges early in development and predicts children’s later vocabulary size and their learning about novel objects (Mundy et al., 2007; Striano, Chen, Cleveland, & Bradshaw, 2006). These findings, among many others, converge to suggest that social interaction plays an important role in early language learning.

If joint understanding and learning develop through social interactions, what are the neural mechanisms that support this transfer of representations? Recent neuroimaging work, primarily with adult participants, has begun to investigate the neural mechanisms that underlie individuals’ representations of shared input. A number of functional magnetic resonance imaging (fMRI) and functional near-infrared spectroscopy (fNIRS) studies have found that adults’ brains exhibit synchronized activity when exposed to the same video or audio recording (Hasson et al., 2004; Wilson, Molnar-Szakacs, & Iacoboni, 2008; Liu et al., 2017). This intersubject temporal correlation of brain activity suggests that individuals exhibit similar neural representations of shared external input. For example, one study reported a relationship between the similarity of participants’ self-reported emotional states during a movie and their neural synchronization in orbital frontal cortex, as well as visual and dorsal attentional networks (intraparietal sulcus and frontal eye field). This suggests that emotional synchrony between individuals was related to similar attentional focus, perception, and understanding of the shared stimulus (Nummenmaa et al., 2012). Kang and Wheatley (2017) reported a similar finding using pupil dilation. Adults listened to a recorded narrative, and pupil size synchrony emerged across participants, especially during emotionally salient periods. These findings suggest that synchronization between adult listeners may be a measure of shared attentional engagement and the emergence of common, dynamic representations when exposed to the same stimulus. Recent developmental research using pupil size synchrony points to similar conclusions (Nencheva et al., 2020).

Converging evidence has demonstrated that intersubject correlation (or neural synchrony) in a network of higher-order brain areas during stories reflects a high-level, joint understanding of narrative information. One set of studies measured coupling between the brain of a speaker telling a personal story and, later, of participants listening to a recording of the story (Iacoboni et al., 2004; Liu et al., 2017; Stephens et al., 2010). Speaker-listener neural coupling was observed not only in areas associated with low-level auditory processing, language production (inferior frontal gyrus), and speech comprehension (superior temporal gyrus and temporo-parietal junction), but also in higher-order extralinguistic, default mode network (DMN) regions, such as the precuneus, parietal lobule, intraparietal sulcus, medial prefrontal cortex, and dorsolateral prefrontal cortex, which are associated with processing of social information (Iacoboni et al., 2004). Importantly, this higher-order neural coupling depends not only on exposure to the same stimulus, but also on a partially or even fully shared understanding of its meaning. For example, in related work with adults who listened to nonsense speech, scrambled speech, or a narrative spoken in a foreign language, synchronization between brains disappeared except in early auditory cortices (Honey et al., 2012; Silbert et al., 2014; Liu et al., 2017). In a recent study of adult-infant coupling during live, face-to-face interactions (playing, singing, and reading), the prefrontal cortex showed the strongest coupling, as well as a relationship to several communicative behaviors (mutual gaze, joint attention to objects, infant emotion, and infant-directed speech) (Piazza et al., 2020).

Given the emerging understanding of brain-to-brain synchrony between people, the next vital question is how it contributes to the real-life goals of communication, such as learning. In particular, is the degree of neural synchrony between a child learner and other young learners predictive of the child’s learning? While previous experiments have largely investigated overall comprehension of a recorded narrative as a measure of the quality of intersubject communication, the present study assessed the correlation between brain-to-brain synchrony and fine-grained word learning from an interactive, naturalistic storybook. Joint book reading in early childhood – a social experience shared by children and caregivers around the world – has been linked to long-term language outcomes, including vocabulary size and literacy (Bus, van IJzendoorn, & Pellegrini, 1995; Dickinson et al., 2012; Noble et al., 2019; Dowdall et al., 2020). We created an original, multifaceted children’s book that enabled analysis of learning at multiple levels, including novel word learning, understanding of object functions, and broader narrative comprehension. Our joint book reading paradigm both afforded temporal control over stimulus presentation and served as a naturalistic context for communication.

To investigate the neural underpinnings of learning from a real-life interaction in preschool-aged children, we used functional near-infrared spectroscopy (fNIRS) to record the neural activity of child participants during joint book reading with an adult experimenter. fNIRS is a neuroimaging modality that uses near-infrared light to record changes in the concentrations of oxygenated and deoxygenated hemoglobin to approximate neural activity. fNIRS is minimally sensitive to motion, which, in addition to permitting naturalistic, reciprocal, real-time interaction, makes it ideal for studies involving children. Following the storybook, we assessed their word learning and story comprehension.

Based on previous adult fMRI and fNIRS studies reporting shared neural representations of stimuli between participants, measured using intersubject correlation or ISC (Hasson et al., 2004; Wilson et al., 2008; Liu et al., 2017), we expected to find significant temporal synchronization of brain activity between child participants during joint book reading with an adult. By reducing individual-specific noise, ISC isolates stimulus-specific neural signals that are consistent across subjects. If an individual child’s neural signature is closer to an overall stimulus-driven pattern, one might expect the child to more effectively extract structure and learn; conversely, if an individual child’s neural signature is less similar to the group as a whole, one might expect the child to show either reduced learning or idiosyncratic patterns of learning. Thus, based on findings that synchronization of neural activity between viewers of a stimulus reflects joint attention and shared understanding (Nummenmaa et al., 2012), we expected that there would be a correlation between child-child neural synchrony in the prefrontal and parietal cortices and learning from the story. Specifically, we anticipated that children whose patterns of neural activity were more (versus less) synchronized with those of other participants would show increased learning of words and other story details. If child-child neural synchrony does predict successful learning, this would enhance our understanding of how the brain engages with the moment-to-moment dynamics of linguistic and social input.

## Method

### Participants

Sixty-nine children aged 3.5-4.5 years old (M = 47.0 months; SD = 3.55 months; 33 females) participated in the study. All children were born full-term, had no history of hearing problems or known developmental delays, and were raised in English-speaking environments (English spoken at least 85% of the time). One adult female experimenter read the story stimulus to all study participants. Five participants were excluded because they refused or were unable to wear the fNIRS cap, one was excluded for moving the cap during the experiment, one was excluded for poor signal quality. An additional seven participants were excluded due to experimenter error or equipment malfunction, and ten were excluded because of an error in the fNIRS equipment configuration (which nullified the data across ten consecutive sessions). The remaining forty-five participants (*M* = 47.5 months; *SD* = 3.68 months; 19 females) were included in analyses.

### Procedure and Stimuli

Each child participant was seated side-by-side with an adult experimenter, who read a digital storybook for 4 minutes and 30 seconds. We designed a custom digital storybook to teach children four novel object-label mappings and expose them to an engaging narrative (Figure 1A). The objects and their labels were selected from the Novel Object and Unusual Name (NOUN) database (Horst & Hout, 2016). The selected objects were highly unlikely to be familiar to participants and had similar visual salience, and their labels (e.g., *foom*, *teebu, glark*, and *koba*) were simple pseudowords, rather than unfamiliar real words, to ensure that participants had no prior exposure to target words. To account for possible subtle differences in label and object salience, children were randomly assigned to see one of two versions of the book; the text was identical in each version, but object-label pairings and order of object presentation were counterbalanced across participants.

**Figure 1.**
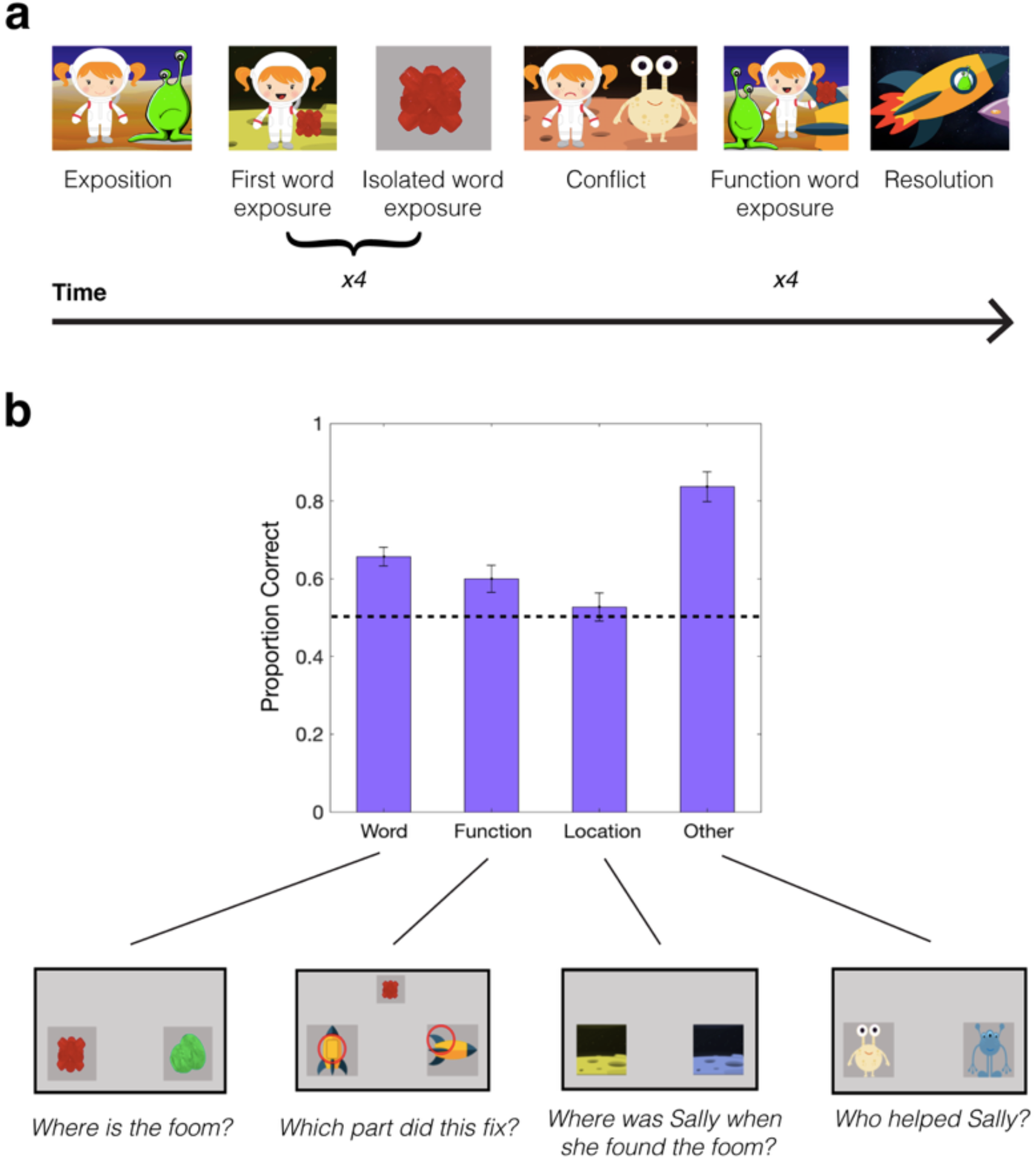
Story stimulus and learning assessment. (a) Schematic timeline of the storybook, which followed a general Western plot structure including exposition, conflict, and resolution. (b) Learning assessment performance, broken down by question category, with an example question from each category. Questions were presented via prerecorded audio clips spoken by the experimenter.

The story adhered to a canonical Western plot structure including exposition, conflict, and resolution. Specifically, it followed a simple narrative of an astronaut traveling through space to find four objects that were needed to fix a broken rocket (Figure 1A). All objects were named three times during the story: once with a colorful backdrop, once in isolation with a black backdrop, and once later in the story with an assigned function (e.g., fixing the engine of the rocket). The story was displayed on a wall-mounted monitor between the child and adult in order to standardize image presentation between the reading and test phases across sessions, and “page-turns” occurred automatically after a predetermined amount of time (approximately 10-15 seconds per page), which controlled for visual exposure to each object. The text of the story appeared on each page, as in typical storybooks (see Supplementary Information for more details), and the experimenter read the text naturally within the time frame of each page. The timing of each page was determined prior to the initiation of data collection based on how long it took the experimenter to read the text. Although advised to adhere strictly to the story’s text, the adult was free to engage with the child by initiating and returning eye contact, nodding, and smiling, as she would during natural book reading. The child participants were able to act naturally, and they occasionally smiled, made eye contact, or vocalized.

### Learning Assessment

Following the story, children participated in a two-alternative forced-choice learning assessment to measure their novel word learning and story comprehension (Figure 1B). Each question appeared on the wall-mounted touch screen monitor, and children were asked via prerecorded audio (spoken by the same adult experimenter who read the story) to select one of the two presented images. Questions were designed to assess receptive learning of novel word labels (e.g., *Where is the foom?, Where is the teebu?*), object functions (*Which part did this fix?*), the locations in which events occurred *(Where was Sally when she found the glark?, Where was Sally when she got lost?*), and general story comprehension (*Who helped Sally?*). Participants were asked a total of 31 questions: 16 to assess novel word learning (four questions for each of the four labeled objects) and 15 to assess story comprehension and understanding of object functions (4 function questions, 5 location questions, and 6 general comprehension). The side of target object presentation was counterbalanced to control for possible side bias. Additionally, the order of the questions was inverted for half of the subjects to minimize the effects of distraction or disinterest near the end of the task. We conducted analyses to ensure that there were no effects of object labeling or side bias on performance (see Supplementary Information).

### Neural Recording

We recorded children’s brain activity during book reading using a LabNIRS system (Shimadzu Scientific Instruments, Columbia, MD). The fNIRS cap covered prefrontal cortex and parts of the temporal and parietal cortex, which have known roles in speech comprehension and social cognition (Wilson et al., 2008). The cap had 20 emitters and detectors, corresponding to a total of 53 channels. We originally planned to measure the neural activity of the adult experimenter, but the quality of the signal from the adult cap declined over the course of data collection (see Supplementary Figure 1), possibly due to a change in elasticity and resulting fit over time. We attempted to reconstruct a single adult brain template by combining high-quality channels from the adult across sessions, but this proved to be underpowered for comparison to data from each child’s brain. Future studies will further explore links between children’s learning and real-time neural coupling between children and adults.

## Results

### Data Preprocessing and Analysis

We conducted analyses to ensure that behavioral results were not biased by order effects, side bias, or object preferences. These analyses (which are described in more detail in the Supplementary Information) resulted in excluding no participants from the main analyses.

We removed motion artifacts using moving standard deviation and spline interpolation (Scholkmann, Spichtig, Muehlemann, & Wolf, 2010) and low-pass-filtered (0.5 Hz) and high-pass filtered (0.02 Hz) the signal to remove physiological noise and signal drift, respectively. To eliminate excessive signal noise, we excluded individual channels in which the time series corresponding to relative concentrations of oxygenated (HbO) and deoxygenated (HbR) were correlated, based on a method established in previous work (Cui et al., 2010). After z-scoring the signal over the duration of joint book reading, we split the time series (4500 data points) into 45 bins and computed windowed Pearson correlations between the concentration changes for each channel. Channels that showed a positive correlation (r ≥ 0) between the HbO and HbR concentration changes averaged across all time bins were excluded. For subsequent analyses, we used the relative concentrations of HbO in each channel to calculate neural synchrony, as recent fNIRS research using naturalistic story stimuli found greater correlations between fMRI BOLD response and HbO concentration changes than between the BOLD and HbR signals (Liu et al., 2017).

Analyses included 53 channels from each child’s fNIRS cap, grouped into three regions of interest (ROIs): prefrontal cortex (PFC) (7 channels), parietal cortex (22 channels), and bilateral temporal cortex (24 channels). We first averaged the HbO signal across channels within each ROI in a given participant. Then, to compute intersubject correlation (ISC), or the degree of synchrony between the participant and all other child participants, we computed a Pearson correlation between the averaged ROI time series (e.g., prefrontal) for the child participant and the average time series across all other child participants in the corresponding ROI. The correlations between learning and neural synchrony were computed with a between-subject Pearson correlation between response accuracy and ISC for each ROI.

### Finding 1: Preschoolers can learn a range of semantic information from a natural storybook

The story was designed to expose children to semantic information at multiple levels of complexity, from the meanings of individual object labels to higher-order, long-timescale narrative information about character goals and conflict resolution. Using a 2AFC task on a touch screen directly following the story, we measured children’s learning of word-object mappings (object labels), the functions of the objects, locations in which the objects were found, and whether certain characters were helpful to the protagonist (Figure 1B). We found that overall learning (collapsed across all question types) was significantly above chance (*t*(44) = 10.6, *p* < .001). More specifically, there was significantly above-chance learning for individual object labels (*t*(44) = 6.5, *p* < .001), object functions (*t*(44) = 2.86, *p* < .01), and other general questions about the story (e.g., the names of main characters; which characters helped the protagonist; *t*(44) = 16.2, *p* < .001). However, preschoolers were unable to recall the locations in which individual objects were found (*t*(44) = 0.759, *p* = .45), likely because the locations (planets) were background scenes that only varied by color.

### Finding 2: Child-child neural synchrony in parietal cortex predicts word learning as well as overall learning across question types

We observed variation in learning both across individual children and across question categories, which enabled us to correlate children’s behavioral performance with their neural synchrony.

We iteratively computed the ISC between each child and the average signal from the other 44 participants in each of three cortical ROIs: frontal, parietal, and temporal (see Methods). For each child, we computed the correlation between this ISC value and their learning of novel words from the story (Figure 2A). This correlation was significant in the parietal ROI (*r*(43) = .33, *p* < .05) but not the other two ROIs (frontal: *r*(43) = .14, *p* = .35; temporal: *r*(43) = .04, *p* = .81). The same pattern was true when we examined the relationship between ISC and overall learning score (accuracy collapsed across all questions; Figure 2B): parietal ROI (*r*(43) = .31, *p* < .05); frontal ROI (*r*(43) = .04, *p* = .78; temporal ROI (*r*(43) = .01, *p* = .99). Correlations between ISC and other forms of learning were not statistically significant [Function: frontal (*r*(43) = −.18, *p* = .24), parietal (*r*(43) = −.08, *p* = .61), temporal (*r*(43) = .04, *p* = .81). Location: frontal (*r*(43) = .02, *p* = .9), parietal (*r*(43) = .07, *p* = .66), temporal (*r*(43) = −.2, *p* = .19). Basic story content: frontal (*r*(43) = −.07, *p* = .6), parietal (*r*(43) = .24, *p* = .12), temporal (*r* = .07, *p* = .65)]. These non-significant findings may be due to the focus of our research question on novel word learning, such that the test phase included more questions about novel word learning than other individual types of questions (by a factor of four).

**Figure 2.**
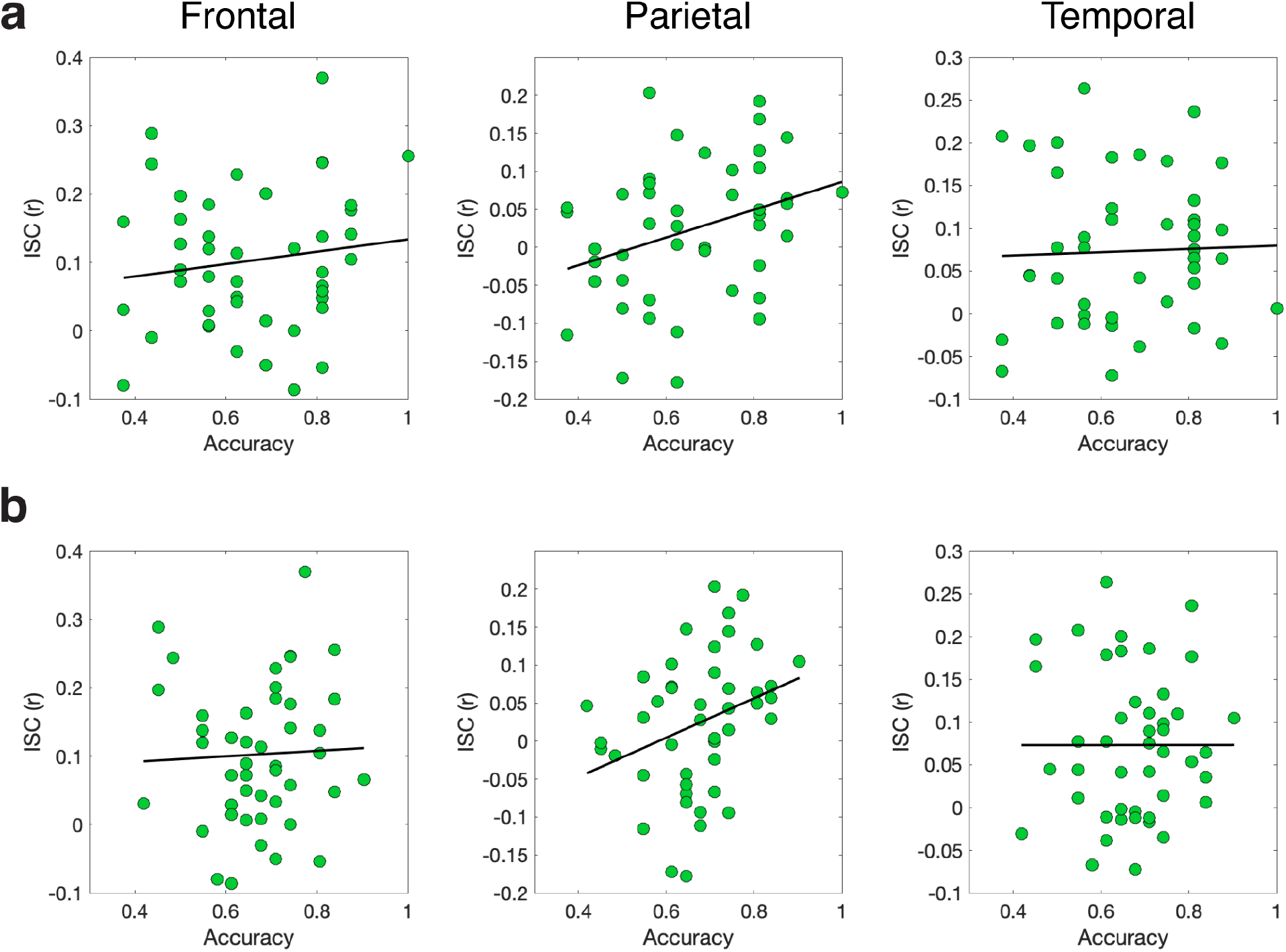
Relationship between intersubject correlation (ISC) and learning of (a) novel words from the story (object label test questions only) and (b) overall story content (collapsed across all test question categories) in each of 3 neural ROIs. N = 45.

### Finding 3: The best overall learners have higher overall ISC

Finally, we performed a median-split analysis by dividing participants into two groups – low-performing and high-performing – based on their learning of novel words and overall story content and comparing the ISC values of these two groups in each ROI (Figure 3). When we split the groups based on novel word learning performance (Figure 3A), a two-way, mixed-effects ANOVA yielded a significant main effect of ROI (*F*(2) = 11.30, *p* < .0001) and a marginally significant main effect of performance group (*F*(1) = 2.95, *p* = .09), but no significant interaction between group and ROI (*F*(2) = 1.14, *p* = .32). Post-hoc independent samples *t*-tests revealed significantly higher ISC in the high-performing (vs. low-performing) group in parietal cortex (one-tailed *t*-test; *t*(42) = 2.45, *p* < .01), but not the other two cortical areas (frontal: *t*(42) = .39, *p* = .35; temporal: *t*(42) = .89, *p* = .19). When we split the groups based on overall story learning performance (i.e., collapsed across all question types; Figure 3B), a mixed effects ANOVA yielded significant main effects of ROI (*F*(2) = 10.18, *p* < .001) and performance group (*F*(1) = 6.05, *p* < .05), but no significant interaction between group and ROI (*F*(2) = 1.8, *p* = .17). Post-hoc independent samples *t*-tests showed significantly higher ISC in the high-performing (vs. low-performing) group in parietal cortex (*t*(42) = 3.02, *p* < .01), marginally higher ISC in frontal cortex (frontal: *t*(42) = 1.63, *p* = .06), and no significant difference between groups in temporal cortex (*t*(42) = .34, *p* = .37).

**Figure 3.**
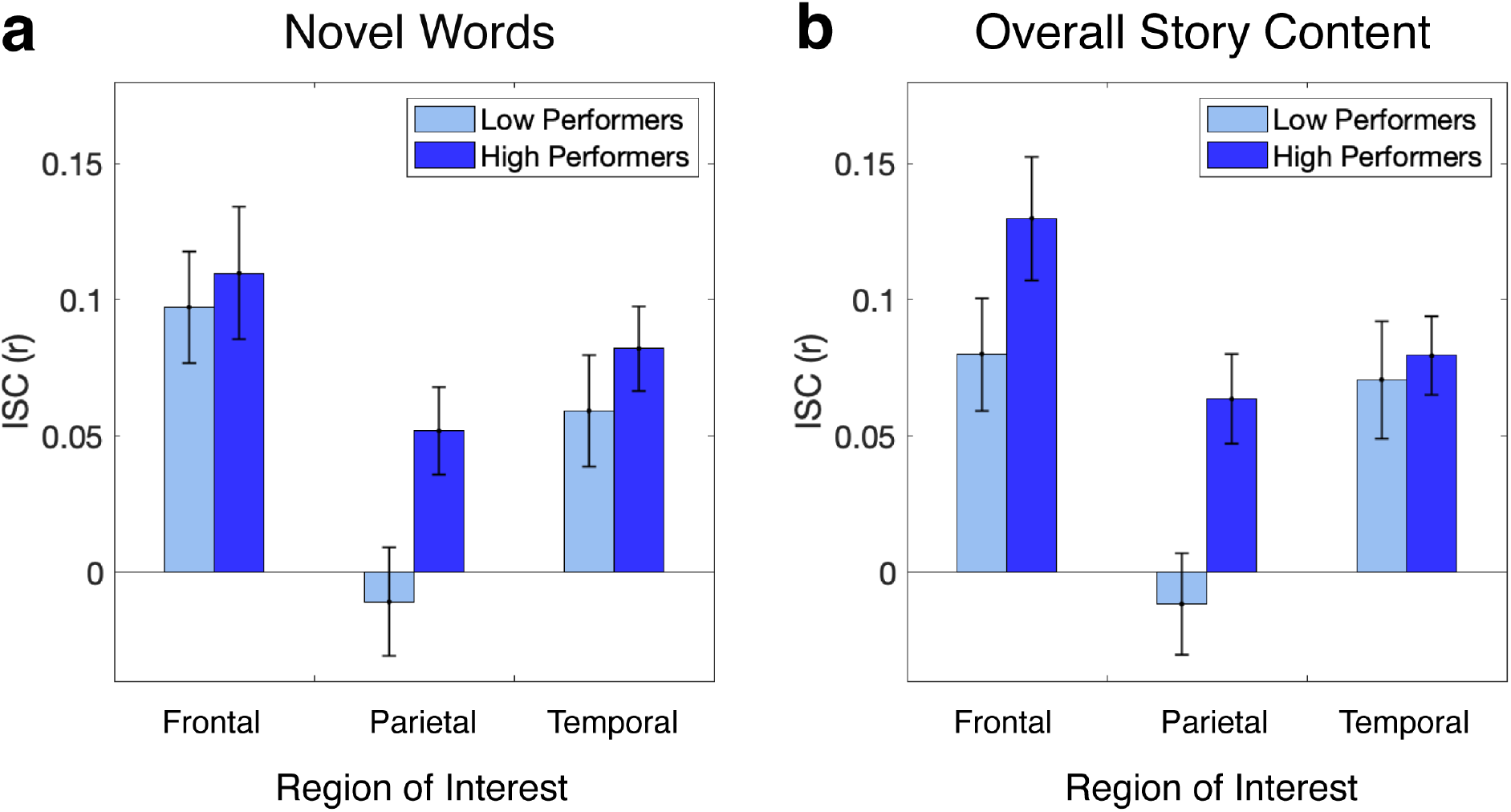
Intersubject correlation (ISC), split into low- and high-performing groups according to learning of (a) novel words and (b) overall story content. Error bars are s.e.m. N = 22 (per performance group).

## Discussion

This study examined the relationship between neural synchrony within a sample of preschool-aged children (during naturalistic, joint book reading with an adult) and learning. The primary goal was to understand if patterns of intersubject neural synchrony would predict young children’s learning of novel words. Broadly, children demonstrated the ability to learn information of varying levels of complexity from the book, including individual object labels, the functions of those objects within the story, and basic character information. We found that neural synchrony (measured with intersubject correlation) between children in parietal channels was significantly correlated with children’s learning, and this was true both for learning of individual words as well as overall learning of story content, collapsed across question types. Furthermore, when we divided participants into groups based on their performance on the comprehension test, we found that higher overall ISC across regions-of-interest was related to children’s higher overall learning from the story. Each child’s ISC provides a proxy for their neural alignment to a signature pattern of processing the story; we found that the closer each child’s neural time series was to this signature pattern (especially in parietal cortex), the better they learned new words introduced throughout the story. This study provides the first evidence of synchronized neural activity between children during joint attention with an adult in the context of a naturalistic interaction, and also of the relationship between this neural synchrony and real-time learning.

Our findings build on previous work demonstrating that the brains of adult observers become coupled in higher-order, default mode network areas (including parietal lobule and medial prefrontal cortex; Iacoboni et al., 2004) when they share a high-level understanding of a stimulus. When high-level understanding breaks down due to scrambling natural speech (Lerner et al., 2011) or translating it into a foreign language (Honey et al., 2012), lower-order sensory areas such as auditory cortex remain synchronized across participants, but synchrony in higher-order areas disappears, revealing that synchrony between participants is driven by more than simple exposure to the same input. Instead, it is based on joint engagement with semantic or narrative-level content. This previous adult work has consistently shown the involvement of both prefrontal and parietal regions in listeners’ high-level interpretations of natural stories (Iacoboni et al., 2004; Stephens et al., 2010; Liu et al., 2017; Yeshurun et al., 2017). In contrast to these studies, which investigated the relationship between neural synchrony among adults and their processing of the “gist” or overall interpretation of a story, our study took the new approach of measuring synchrony between children and focusing on its relationship to novel word learning. We found that this relationship was significant only in parietal cortex, and the lack of an effect in prefrontal cortex could be due to several factors. For instance, it is possible that in early childhood, parietal brain areas play a strong role in tracking narrative content, whereas the prefrontal cortex does not become functionally connected to the rest of the default mode network (as characterized in adults in the studies above) until later in development. Alternatively, the process of tracking individual words’ meanings may be more strongly linked to attentional factors regulated by parietal cortex, in which case we would predict to see a similar pattern of results if we tested novel word learning in adults. Interestingly, when we divided our participants into groups based on their overall learning across all question types (which is analogous to the comprehension tests in previous adult studies), we did find differences between high- and low-performing groups in both parietal *and* frontal areas. Future research directly comparing the role of neural synchrony in learning multiple levels of information from stories—isolated novel object labels and their functions, character motives, and overall narrative arcs—in both adults and children will help clarify the mechanistic role of multiple brain areas in the moment-to-moment encoding, maintenance, and recall of natural input.

This work sheds new light on the neural underpinnings of early word learning. Previous electrophysiological and neuroimaging studies have tracked dynamic changes in neural activation patterns in the hippocampus, left temporal lobe, and inferior frontal gyrus during word learning in adults (Shtyrov, 2012). However, young children’s word learning is subject to a myriad of unique factors that distinguish this process from adult word learning: when a preschooler hears a new word within a story, they are still learning how to integrate words within the context of long-timescale narrative arcs, and when an infant hears their parent say a new word, they are still learning to segment acoustic input into syllables. Thus, the aggregate of developmental processes impacting the learning of an individual word should be carefully considered in studies of language development, and approaches comparing child-child synchrony in multiple brain areas and across developmental stages will be particularly useful in this effort. Our finding that neural synchrony also predicted learning of longer-timescale information beyond word meaning (e.g., which characters were helpful) could be expanded upon in future studies investigating how structural disruptions of stories (e.g., via scrambling; Lerner et al., 2011) impact both neural synchrony in different brain areas and comprehension.

Our study used a joint book reading paradigm, and this social form of engagement with language has been shown to benefit learning. While children have the ability to learn new words through passive listening to storytelling (Elley, 1989), studies of preschool-aged children suggest that interactive, shared reading facilitates greater vocabulary acquisition than passive listening (Whitehurst et al., 1988; Hargrave & Sénéchal, 2000; Opel, Ameer, & Aboud, 2009). Analyses of fully natural infant-adult shared reading experiences reveal that these interactions take on a dialogic, contingent structure, with adult communication tailored to children’s developmental level and real-time feedback (Ninio & Bruner, 1978; Vygotsky, 1978; Dowdall et al., 2020). To do so, caregivers adopt many behaviors that engage children and enhance word learning during book reading, such as recasting and repeating unfamiliar words, changing the content of their speech, asking ‘what’ and ‘where’ questions, and connecting story content to the child’s personal experiences (Ard & Beverly, 2004; Hayden & Fagan, 1987; Ninio, 1980; Wheeler, 1983). Although our design was naturalistic and allowed for joint engagement between the experimenter and child, our use of one experimenter and timed page-turns limited reader-child interaction to some degree. In the future, assessing how social engagement and behavioral feedback are reflected in children’s neural representations during dialogic book reading may help us understand why this type of interaction positively affects language outcomes. Incorporating children’s own caregivers into future fNIRS studies will provide insights into the neural mechanisms supporting parents’ tailoring of behavioral cues to promote joint attention.

Although technical limitations prevented us from analyzing the role of adult-child coupling in learning (see Supplementary Figure 1), this is an exciting area for future research. Recent EEG (Pan et al., 2020) and fMRI (Nguyen et al., 2020; Meshulam et al., 2020) studies of adults have begun to explore the relationship between teacher-student coupling and learning outcomes, but the arena of early, interactive learning from caregivers is underexplored. Future research could investigate the role of different behavioral cues (prosody, eye gaze, gesture) at key moments in a story, such as before or after initial exposure to a novel word, in aligning children’s neural representations of semantic content with the adult storyteller’s representations. Additionally, although our analyses focused on mirrored, one-to-one synchrony between the child listeners’ brains, adult-child interactions are more likely characterized by non-mirrored coupling functions (Hasson & Frith, 2016). For example, leader-follower relationships (which might occur when an adult has prior knowledge that enables faster anticipation of plot development compared to a child) could be evaluated using lagged neural synchrony measures. In a more improvisational, joint storytelling context (such as during free play), the two brains may synergistically constrain or adapt to the each other to invent new content together. To further understand the link between adult-child neural coupling and word learning, it will be fruitful to track how a wide range of coupling patterns maps onto different developmental stages and interaction contexts.

## Acknowledgements

We thank the participating families and the members of the Princeton Baby Lab. This work was supported by grants from the National Institute of Child Health and Human Development to C.L.W. (R01HD095912, R03HD079779) and a Princeton University C. V. Starr Fellowship to E.A.P.

## Notes

### Competing Interest Statement

The authors have declared no competing interest.

