## Supplementary Information for "Neural synchrony predicts children’s learning of novel words"

*Story Design*

We designed a storybook to teach children novel words and expose them to a narrative (Figure 1A), their comprehension of which we later assessed to evaluate learning. The story integrates four objects into a narrative designed to engage participants. Objects to which children are unlikely to have had previous exposure were selected from the Novel Object and Unusual Name (NOUN) database and other online sources (Horst & Hout, 2016). The selected objects have similar brightness and complexity to avoid differential object salience. We used pseudowords (e.g., “foom” and “teebu”) obtained from the NOUN Database, rather than unfamiliar real words, to ensure that participants had no prior exposure to target words and that existing vocabulary did not confound word learning. Object labels were selected to be relatively simple (containing one or two syllables) and consistent with English phonology to facilitate learning. To account for subtle differences in label and object salience, we presented two versions of the story. The text was identical in both versions of the story, but the object-label pairings and the order of object presentation were distinct.

The story follows a simple narrative of an astronaut traveling through space to find four objects to fix her friend’s rocket. All objects are named three times throughout the story: twice at their initial presentation and once when they are assigned a function (e.g., fixing a specific part of the rocket). The story was displayed on a wall-mounted monitor to standardize image presentation between the reading and test phases. “Page-turns” were automated to control participant exposure to each object.

The study was designed to assess participants’ word learning based on auditory object labeling during book reading. To replicate a naturalistic shared book reading experience between an adult and child, story text was included on the stimulus, because removing the story’s text would reduce the similarity between this paradigm and its real-life counterpart. Displaying text also ensured that the adult experimenter exposed each child to precisely the same story content.

Furthermore, an eye-tracking study found that four- and five-year-old children (most of whom were older than the participants in the present study) spend minimal time (2.7% of total looking time) fixating on the regions of the page containing text (Justice et al., 2005). Because some children in the study’s age range are able to read simple words, such as the object labels employed in the study (*foom, glark, teebu,* and *koba*), we asked participants’ parents to complete a questionnaire about their children’s reading ability. Only two parents reported that their children were able to read. Importantly, no parents reported that their children could read unfamiliar words such as “blazer” or “spade,” which were similar in complexity to the novel words presented in the story.

*Learning Assessment*

Prior to data analysis, we analyzed children’s response data for potential participant exclusion. To verify that there was no object preference, bias in object labeling, or order effect of object presentation, we conducted a one-way ANOVA of response accuracy for objects by image and another by label, and we found no significant difference for questions by image ($F(3,176)$ =

0.16, $p = 0.92$) or by object label ($F(3,180) = 2.07$, $p = 0.11$). Finally, we established a threshold of 85% percent to test for side bias and found that no subjects exceeded this threshold.

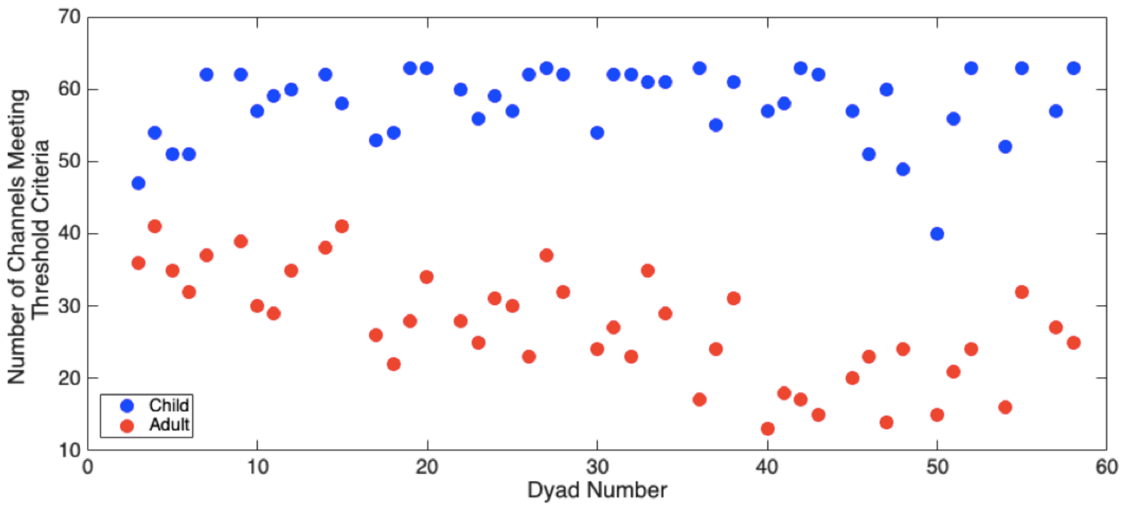


**Supplementary Fig. 1**. Number of channels meeting criteria for acceptable signal quality across dyads, in the children (blue) and adult (red).
